# Association between breastfeeding and DNA methylation over the life course: findings from the Avon Longitudinal Study of Parents and Children (ALSPAC)

**DOI:** 10.1101/800722

**Authors:** Fernando Pires Hartwig, George Davey Smith, Andrew J Simpkin, Cesar Gomes Victora, Caroline L. Relton, Doretta Caramaschi

**Affiliations:** Postgraduate Programme in Epidemiology, Federal University of Pelotas, Pelotas, Brazil; MRC Integrative Epidemiology Unit, University of Bristol, Population Health Science, Bristol Medical School, Bristol, United Kingdom; Insight Centre for Data Analytics, National University of Ireland, Galway, Ireland

**Keywords:** Breastfeeding, Life-course, DNA methylation, Epigenome-wide association study

## Abstract

**Background:** Breastfeeding is associated with short and long-term health benefits. Long-term effects might be mediated by epigenetic mechanisms, yet a recent systematic review indicated that the literature on this topic is scarce. We performed the first epigenome-wide association study of infant feeding, comparing breastfed vs non-breastfed children. We measured DNA methylation in children from peripheral blood collected in childhood (age 7, N=640) and adolescence (age 15-17, N=709) within the Accessible Resource for Integrated Epigenomic Studies (ARIES) project, part of the larger Avon Longitudinal Study of Parents and Children (ALSPAC) cohort. Cord blood methylation (N=702) was used as a negative control for potential pre-natal residual confounding.

**Results:** Two differentially-methylated sites presented directionally-consistent associations with breastfeeding at ages 7 and 15-17, but not at birth. Twelve differentially-methylated regions in relation to breastfeeding were identified, and for three of them there was evidence of directional concordance between ages 7 and 15-17, but not between birth and age 7.

**Conclusions:** Our findings indicate that DNA methylation in childhood and adolescence may be predicted by breastfeeding, but further studies with sufficiently large samples for replication are required to identify robust associations.

## Background

Breastfeeding has clear short-term health benefits, particularly in reducing the risk of infections in childhood. Accumulating evidence indicates that breastfeeding may also have long-term effects on health outcomes and human capital, as well as benefit maternal health [1]. For example, being breastfed has been associated with better performance in intelligence quotient (IQ) tests in a meta-analysis based on a systematic literature review [2], in population-based birth cohorts with different confounding structures [3], and in the single large randomized controlled trial on this subject [4].

The mechanisms underlying the long-term effects of breastfeeding are not fully understood. Such mechanisms clearly must persist over time after weaning – in other words, become “imprinted” in the organism [5]. In the case of other early-life exposures such as maternal smoking during pregnancy, there is evidence of long-term associations with offspring DNA methylation [6] – i.e., addition of a methyl (–CH_3_) group to DNA at the 5’ position of a cytosine base, typically in cytosine-guanine (CpG) dinucleotides located in DNA sequences called CpG islands, which are rich in CpG dinucleotides [7, 8]. DNA methylation is one type of a broader class of biological processes known as epigenetics, which encompasses mitotically heritable events – other than changes in the DNA sequence itself – involved in gene expression regulation. Epigenetic processes play a key role in developmental processes [9, 10], and have more recently been linked to disease processes [11, 12, 13, 14], although causal claims have been overstated in many cases [15].

Some evidence suggests that breastfeeding might influence DNA methylation through the effects of some of its nutritional components [16] or through the microbiome, which is shaped by early feeding habits [17]. However, according to a recent systematic literature review [18], the overall evidence on the potential effects of breastfeeding on DNA methylation is scarce. Our aim was to perform a genome-wide assessment of the association between breastfeeding and DNA methylation in childhood, characterise – if present – the pattern of this association and investigate whether it persists until adolescence in a population-based study in England.

## Methods

### Study setting and participants

Study subjects were part of the Accessible Resource for Integrated Epigenomic Studies (ARIES) [19], a sub-sample of the Avon Longitudinal Study of Parents and Children (ALSPAC) for which methylation data were collected. ALSPAC is a population-based, prospective birth cohort of women and their children [20, 21, 22]. All pregnant women living in the geographical area of Avon (UK) with expected delivery date between 1 April 1991 and 31 December 1992 were invited to participate. Approximately 85% of the eligible population was enrolled, totalling 14,541 pregnant women who gave informed and written consent. Information on the data collection and availability can be found at http://www.bris.ac.uk/alspac/researchers/data-access/data-dictionary/. Ethical approval for the study was obtained from the ALSPAC Ethics and Law Committee and the Local Research Ethics Committees.

Our analysis was focused on the offspring born in 1991-1992. The analyses were restricted to singletons or only to one participant out of a twin pair, selected at random. Individuals with missing information for the exposure, outcome or covariates (described below) were excluded.

### Study variables

#### DNA methylation

DNA methylation in white blood cells was measured in ARIES offspring at three time points: at birth (cord blood), and at 7 and 15-17 years of age (peripheral blood). DNA samples underwent bisulphite conversion using the Zymo EZ DNA methylationTM kit (Zymo, Irvine, CA). The Illumina HumanMethylation450 BeadChip was used for genome-wide epigenotyping. The arrays were scanned using an Illumina iScan, and initial quality checks performed using GenomeStudio version 2011.1. We excluded single nucleotide polymorphisms, probes with a high detection P-value (ie, P-value>0.05 in more than 5% samples) and sex chromosomes. Methylation data normalisation was carried out using the “Tost” algorithm to minimise non-biological between-probe differences [23], as implemented in the “watermelon” R package [24]. All processing steps used the “meffil” R package[25].

The outcome variables of this study were cord and peripheral blood (ages 7 and 15) DNA methylation levels in ∼470,000 CpG sites. Methylation was analysed as beta values, which vary from 0 to 1 and indicate the proportion of cells methylated at a particular CpG [26]. Regression coefficients and standard errors were multiplied by 100, so that they can be interpreted as percent point differences in average DNA methylation at a given CpG site.

#### Breastfeeding

Breastfeeding data were collected through questionnaires answered by the mothers when their offspring were (on average) four weeks, six months and 15 months old, and combined into four different breastfeeding categorisations:

- A binary indicator of whether the individual was ever breasted (regardless of duration).
- Breastfeeding duration groups, defined as follows: 0=never breastfed; 1=1 day to 3 months of duration; 2=3.01 to 6 months; 3=6.01 to 12 months; and 4=more than 12 months.
- Same as the above, but coding each category as a number and treating this as a continuous variable, thus assuming a linear trend per unit increase in duration category.
- Breastfeeding duration in months, as a continuous variable, thus assuming a lineartrend per month increase in breastfeeding duration.

#### Covariates

Covariates were selected mostly based on a conceptual model encoded in the form of a directed acyclic graph (DAG) that we defined previously [18]. The following covariates were used:

- Sociodemographic: an indicator of whether the participant had white ethnic background (informed by mothers at 32 weeks of gestation), and the top two ancestry-informative principal components estimated using genome-wide genotyping data [27].
- Family socioeconomic position: to avoid collinearity issues, we used only the mother’s highest educational qualification (informed by the mothers themselves at 32 weeks of gestation).
- Maternal characteristics: parity (informed by the mothers at 18 weeks of gestation), height, pre-pregnancy weight (informed by the mothers themselves at 12 weeks of gestation), age at birth (calculated from mother’s date of birth and date of delivery) and folic acid supplementation (informed by the mothers at 18 and 32 weeks of gestation).
- Gestational characteristics: maternal smoking during pregnancy (informed by the mothers at 18 weeks of gestation), type of delivery (informed by the mothers when their offspring were eight weeks old), gestational age (calculated from the date of the mother’s last menstrual period reported at enrolment; when the mother was uncertain of this or when it conflicted with clinical assessment, the ultrasound assessment was used; where maternal report and ultrasound assessment conflicted, an experienced obstetrician reviewed clinical records and provided an estimate) and birthweight (from obstetric data, measures from the ALSPAC team and notifications or clinical records).

Although not included in the DAG, participant’s sex and age at blood collection were also selected as covariates. Given that they are associated with DNA methylation but are not influenced by breastfeeding, adjusting for those two covariates may improve power by reducing variance in DNA methylation. We also adjusted for estimated cell counts using Bakulski’s [28] (for cord blood) or Houseman’s (for peripheral blood) [29] methods to account for methylation differences due to cell composition. Finally, a surrogate variable analysis was performed on the methylation data using the “sva” R package, and the surrogate variables not associated with breastfeeding were additionally included as covariates to adjust for batch effects [30].

### Statistical analyses

We conducted an epigenome-wide association study (EWAS) of any reported breastfeeding (including mixed breast- and formula-feeding and in combination with other foods). The main EWAS analyses considered breastfeeding as the exposure in two categorisations: i) none vs. any; ii) duration categories, assuming a linear trend. We opted for this categorisation rather than the continuous breastfeeding variable because the latter is likely prone to substantial measurement error and digit preference in the self-reported months of duration. Moreover, assuming a linear effect over the entire range of breastfeeding duration (which entails assuming, for example that the effect of changing from 0 to 1 month is the same as the effect of changing from 15 to 16 months) seems less plausible then a linear trend over duration categories (which entails assuming, for example, that the effect of changing from 0-3 to 3.01-6 months is the same as the effect of changing from 6.01-12 to >12 months). The outcome was DNA methylation measured at ∼470,000 CpG sites in peripheral blood at the age of 7 years. The association of methylation at CpGs at age 7 with suggestive evidence, here defined as achieving a P-value<5.0×10^−6^, and breastfeeding was further analysed using additional categorisations of theexposure (breastfeeding). We also investigated whether the signal persisted over time by analysing the association of CpG methylation at age 15-17 and breastfeeding (ever vs never and duration categories). Cord blood methylation was analysed as a negative control [31], under the assumption that at least some of possible pre-natal residual confounding would result in associations between breastfeeding and cord blood methylation. Two analysis models were performed on the subjects with complete covariate data available (N=702): i) adjusting only for estimated cell composition and batch effects, and ii) adjusting for all covariates. These models are hereafter referred to as minimally-adjusted and fully-adjusted, respectively. All analyses were performed using heteroskedasticity-consistent standard errors, implemented using the “lmtest”, “MASS” and “sandwich” R packages.

The EWAS results were further used to identify differentially methylated regions (DMRs) in relation to breastfeeding. DMRs were identified using the Comb-P method, which tags regions enriched for low P-values while accounting for auto-correlation and multiple testing [32, 33]. Following the criteria used by Sharp et al. [34], a region was classified as a DMR if: i) it contained at least two CpGs; ii) all CpGs in the region are within 1000 bp of at least another CpG in the same region; and iii) the auto-correlation and multiple-testing corrected (upon applying Stouffer-Liptak-Kechris and Sidak methods, respectively) P-value for the region was <0.05. The CpGs belonging to the identified DMRs were analysed further to assess if breastfeeding had a consistent effect across the DMR (ie, if CpGs in the DMR generally presented greater or lower levels of methylation according to breastfeeding) using linear mixed models to account for the correlation between CpGs assuming that they are nested within individuals. Therefore, each CpG in a given DMR was treated as a repeated measure of DNA methylation, and the regression coefficient indicates the average difference in DNA methylation levels comparing breastfed and never breastfed individuals, averaging across all CpGs in the DMR. This was implemented using the “nlme” R package. This was complemented by evaluating, for each DMR, the directional consistency of each CpG across time points using a sign test. Analyses were performed using R 3.4 (R Core Team (2018). R: A language and environment for statistical computing. R Foundation for Statistical Computing, Vienna, Austria. URL https://www.R-project.org).

## Results

### Description of study participants

Supplementary Table 1 displays the characteristics of the study participants. There were 702 (birth), 640 (age 7) and 709 (age 15-17) individuals with non-missing information for all study variables (corresponding to approximately 70% of all ARIES participants). In general, the subset included in our analysis was similar to the entire ARIES dataset. The largest differences were observed for maternal education at birth (with the mothers of included individuals having slightly higher educational attainment) and ethnicity (with the proportion of individuals of white ethnic background being slightly higher in the included individuals). Previous analyses indicated that ARIES is reasonably representative of the entire ALSPAC cohort [19].

### Association of breastfeeding with single CpG sites

Figures 1 and 2 provide an overall view of the EWAS results. There was no strong indication of genome-wide inflation for breastfeeding analysed in duration categories, assuming a linear trend (genomic inflation factor of 0.97), but there was some indication for the “ever breastfeeding” variable (genomic inflation factor of 1.10). Importantly, the bulk of the distribution closely resembled the expected under the null, with the deviation occurring in the right tail of the distribution of P-values. This may be due to breastfeeding having small effects on DNA methylation (in which case detection would require larger samples) in many regions of the genome, rather than due to the presence of systematic bias in the results.

**Figure 1.**
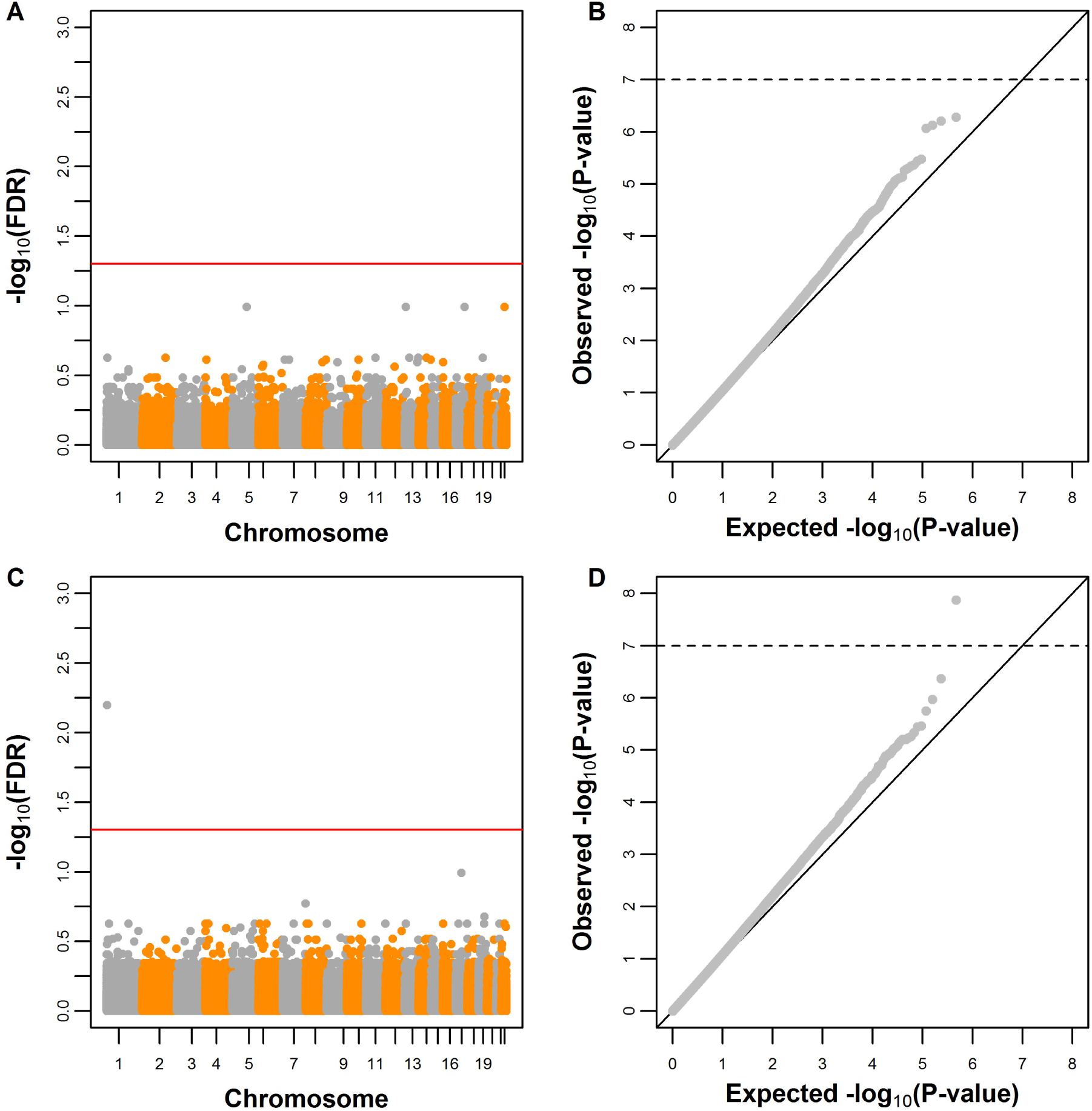
Manhattan and Q-Q plots of the breastfeeding EWAS, comparing peripheral blood methylation at age 7 between never vs. ever breasted individuals. A,C: Manhattan plots. B,D: Q-Q plots. A,B: Minimally-adjusted model. C,D: Fully-adjusted model.

**Figure 2.**
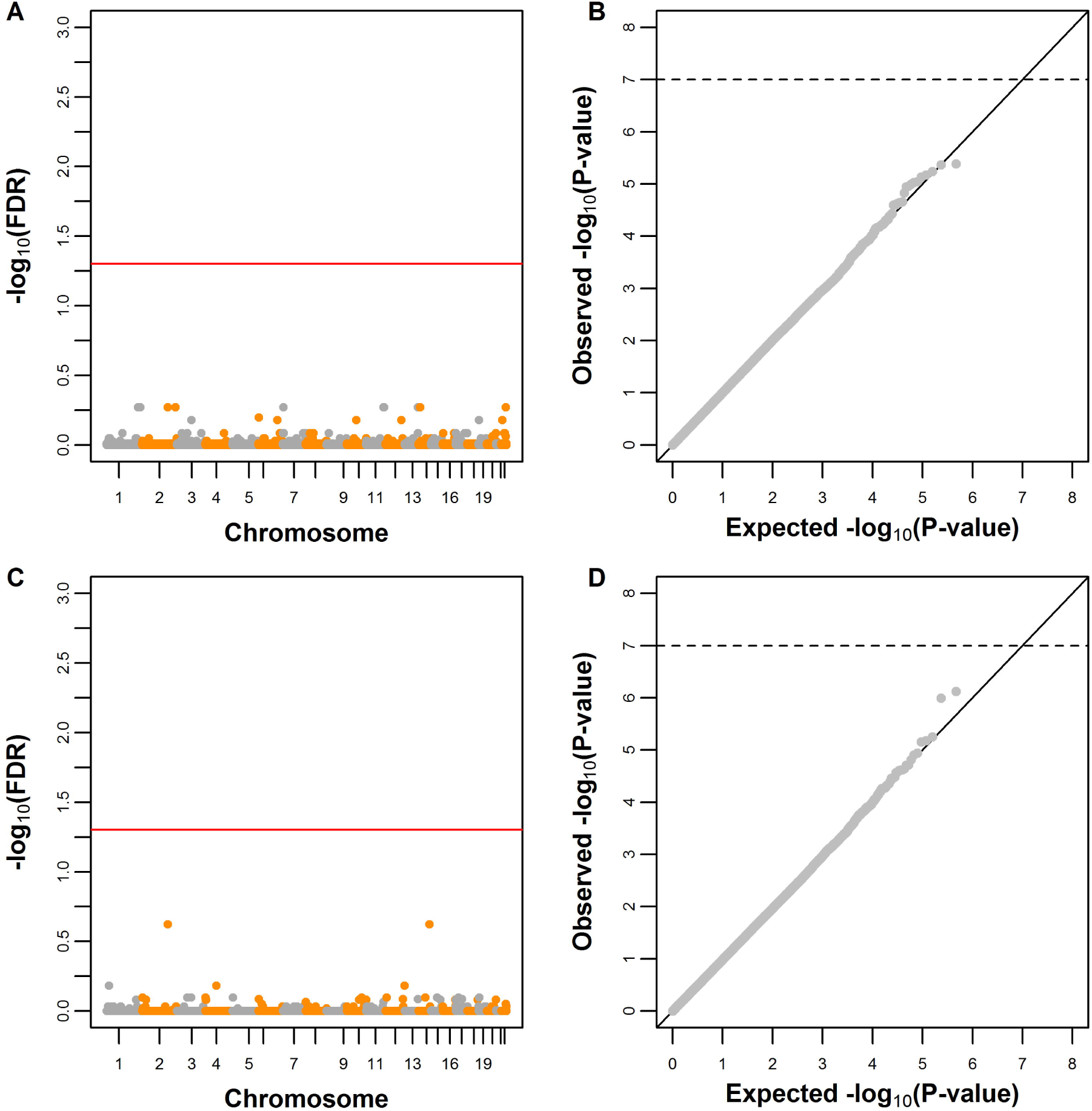
Manhattan and Q-Q plots of the breastfeeding EWAS, comparing peripheral blood methylation at age 7 according to breastfeeding duration (in categories, assuming a linear trend). A,C: Manhattan plots. B,D: Q-Q plots. A,B: Minimally-adjusted model. C,D: Fully-adjusted model.

Regarding ever breastfeeding, no CpGs achieved the conventional threshold of FDR<0.05 (which approximately corresponds to a P-value of 1.0×10^−7^) in the minimally-adjusted model, although a few achieved FDR<0.20 (which approximately corresponds to a P-value of 1.0×10^−6^). In the fully-adjusted model (Table 1), one CpG (cg11414913) achieved FDR<0.05, and there was suggestive evidence of association for six additional sites (cg00234095, cg04722177, cg03945777, cg17052885, cg05800082 and cg24134845; see Supplementary Table 2 for a description of those CpGs). The results for breastfeeding coded as a categorical variable in duration categories (assuming a linear trend) were null, with no CpGs achieving even suggestive levels of association. This suggests that, if breastfeeding is associated with peripheral blood DNA methylation, the association depends more on whether or not the individual was ever breastfed than breastfeeding duration.

**Table 1.** Average percent point differences (β) in DNA methylation at age 7 (N=640) according to breastfeeding.

| Breastfeeding | Statistic | CpG |  |  |  |  |  |  |
| --- | --- | --- | --- | --- | --- | --- | --- | --- |
|  |  | cg11414913 | cg00234095 | cg04722177 | cg03945777 | cg17052885 | cg05800082 | cg24134845 |
| **Binary (ever vs. never)** | P-value | 5.2×10<sup>-8</sup> | 4.9×10<sup>-7</sup> | 2.7×10<sup>-6</sup> | 3.2×10<sup>-6</sup> | 4.9×10<sup>-6</sup> | 5.8×10<sup>-6</sup> | 3.3×10<sup>-5</sup> |
| | $\beta$ (SE) | -3.19 (0.59) | -1.74 (0.35) | -2.90 (0.62) | -0.84 (0.18) | 1.79 (0.39) | 1.05 (0.23) | 0.23 (0.06) |
| **Categories** | P-value | - | - | - | - | - | - | - |
| 0 | $\beta$ (SE) | 0 (Ref.) | 0 (Ref.) | 0 (Ref.) | 0 (Ref.) | 0 (Ref.) | 0 (Ref.) | 0 (Ref.) |
|  | P-value | 1.5×10<sup>-6</sup> | 1.2×10<sup>-7</sup> | 5.3×10<sup>-4</sup> | 2.9×10<sup>-5</sup> | 8.2×10<sup>-6</sup> | 1.7×10<sup>-6</sup> | 6.8×10<sup>-5</sup> |
| 0.01-3 months | $\beta$ (SE) | -3.19 (0.66) | -2.02 (0.38) | -2.45 (0.71) | -0.85 (0.20) | 1.85 (0.41) | 1.19 (0.25) | 0.25 (0.06) |
|  | P-value | 5.4×10<sup>-7</sup> | 3.3×10<sup>-5</sup> | 5.8×10<sup>-5</sup> | 0.005 | 6.8×10<sup>-5</sup> | 6.4×10<sup>-4</sup> | 0.011 |
| 3.01-6 months | $\beta$ (SE) | -3.50 (0.70) | -1.88 (0.45) | -3.22 (0.80) | -0.66 (0.23) | 1.85 (0.47) | 0.94 (0.28) | 0.17 (0.07) |
|  | P-value | 2.5×10<sup>-5</sup> | 3.2×10<sup>-4</sup> | 5.9×10<sup>-5</sup> | 7.4×10<sup>-5</sup> | 6.1×10<sup>-6</sup> | 0.001 | 2.2×10<sup>-4</sup> |
| 6.01-12 months | $\beta$ (SE) | -3.00 (0.71) | -1.59 (0.44) | -3.05 (0.76) | -0.90 (0.23) | 2.02 (0.45) | 0.87 (0.27) | 0.24 (0.06) |
|  | P-value | 5.8×10<sup>-4</sup> | 0.037 | 1.1×10<sup>-6</sup> | 1.2×10<sup>-4</sup> | 0.008 | 0.001 | 4.4×10<sup>-4</sup> |
| >12 months | $\beta$ (SE) | -2.96 (0.86) | -0.93 (0.44) | -3.79 (0.78) | -0.99 (0.26) | 1.29 (0.49) | 1.04 (0.31) | 0.25 (0.07) |
| **Linear trend of categories** | P-value | 0.036 | 0.832 | 1.7×10<sup>-4</sup> | 0.007 | 0.067 | 0.230 | 0.020 |
| | $\beta$ (SE) | -0.42 (0.20) | -0.02 (0.11) | -0.70 (0.19) | -0.16 (0.06) | 0.19 (0.10) | 0.08 (0.07) | 0.04 (0.02) |
| **Continuous (in months)** | P-value | 0.080 | 0.766 | 2.5×10<sup>-4</sup> | 0.035 | 0.966 | 0.399 | 0.289 |
| | $\beta$ (SE) | -0.09 (0.05) | 0.01 (0.03) | -0.18 (0.05) | -0.03 (0.02) | 0.00 (0.03) | 0.01 (0.02) | 0.00 (0.00) |

SE: standard error.

Table 1 shows that methylation in the cg11414913 CpG was 3.2 percent points lower (P=5.2×10^−8^) in ever breastfed children. There was also suggestive evidence for an association between breastfeeding and lower methylation in the cg00234095 (β=-1.7; P=4.9×10^−7^), cg04722177 (β=-2.9; 2.7×10^−6^), and cg03945777 (β=-0.8; P=3.2×10^−6^) sites, and for higher methylation in the cg17052885 (β=1.8; P=4.9×10^−6^), cg05800082 (β=1.1; P=5.8×10^−6^), and cg24134845 (β=0.2; P=3.3×10^−5^) sites. The evidence of an association was greatly attenuated when breastfeeding was analysed continuously (in months), and the regression coefficients were generally similar among different categories of breastfeeding duration. Those results indicate that the association between breastfeeding and peripheral blood DNA methylation is unlikely to follow a dose-response relationship, but presents a threshold (ever vs. never) pattern.

Table 2 displays the association between ever breastfeeding and peripheral blood methylation at different ages in the CpGs identified in the EWAS. The cg11414913 CpG presented a persistent, directionally-consistent association with breastfeeding at the age of 15-17 years (β=-2.8; P=0.004), and no strong evidence of association at birth (β=-0.4; P=0.631). The cg05800082 CpG presented a similar pattern, although the point estimate was attenuated compared to age 7 years, and presented rather weak statistical evidence of association at the age of 15-17 years (β=0.6; P=0.083). However, it was reassuring that its point estimate at birth (β=-0.5; P=0.144) was directionally inconsistent with the results at later ages. The CpGs cg00234095, cg03945777 and cg24134845 presented evidence of association only at age 7, suggesting that their association with breastfeeding does not persist until the ages of 15-17. DNA methylation at birth in the two remaining CpGs was associated with breastfeeding in the same direction as the association at the age of 7, suggesting that those associations are substantially influenced by some unaccounted bias source (e.g., unmeasured confounders).

**Table 2.** Average percent point differences (β) in DNA methylation at different ages according to breastfeeding.

| CpG | Time point | $\beta$ | SE | P-value |
| --- | --- | --- | --- | --- |
| cg11414913 | At birth (N=702) | -0.44 | 0.91 | 0.631 |
| | 7 years (N=640) | -3.19 | 0.59 | $5.2 \times 10^{-8}$ |
|  | 15-17 years (N=709) | -2.47 | 0.85 | 0.004 |
| cg00234095 | At birth (N=702) | 0.59 | 0.57 | 0.296 |
| | 7 years (N=640) | -1.74 | 0.35 | $4.9 \times 10^{-7}$ |
|  | 15-17 years (N=709) | 0.29 | 0.43 | 0.505 |
| cg04722177 | At birth (N=702) | -1.50 | 0.70 | 0.032 |
| | 7 years (N=640) | -2.90 | 0.62 | $2.7 \times 10^{-6}$ |
|  | 15-17 years (N=709) | -1.05 | 0.78 | 0.180 |
| cg03945777 | At birth (N=702) | 0.42 | 0.3 | 0.158 |
| | 7 years (N=640) | -0.84 | 0.18 | $3.2 \times 10^{-6}$ |
|  | 15-17 years (N=709) | 0.10 | 0.29 | 0.742 |
| cg17052885 | At birth (N=702) | 1.32 | 0.57 | 0.022 |
| | 7 years (N=640) | 1.79 | 0.39 | $4.9 \times 10^{-6}$ |
|  | 15-17 years (N=709) | -0.29 | 0.47 | 0.547 |
| cg05800082 | At birth (N=702) | -0.53 | 0.36 | 0.144 |
| | 7 years (N=640) | 1.05 | 0.23 | $5.8 \times 10^{-6}$ |
|  | 15-17 years (N=709) | 0.56 | 0.32 | 0.083 |
| cg24134845 | At birth (N=702) | 0.04 | 0.07 | 0.535 |
| | 7 years (N=640) | 0.23 | 0.06 | $3.3 \times 10^{-5}$ |
|  | 15-17 years (N=709) | 0.00 | 0.08 | 0.991 |

SE: standard error.

### Association between breastfeeding and methylation regions

Given that quantile-quantile plots were suggestive of small effects of breastfeeding on DNA methylation in many regions of the genome, we complemented the ever breastfeeding EWAS with a search for differentially methylated regions (DMRs) – i.e., two or more CpGs enriched for low P-values of the association with breastfeeding (see the Methods for details). 12 DMRs were identified at age 7 (Table 3 and Supplementary Table 3). There was no strong indication that the association of breastfeeding with different CpGs in the same DMR was generally directionally consistent (Table 3). However, regarding directional concordance for each CpG across time points, four DMRs presented evidence of concordance between 7 and 15-17 years, but not between methylation and birth and at age 7: 18:106178-106850, 9:91296-92146, 22:255590-256045, and 8:409905-410098 (Table 4). For two DMRs (5:97867-98797 and 1:425524-426297), there was evidence for directional concordance between birth and 7 years of age, suggesting that the associations between breastfeeding and methylation at age 7 in the CpGs in those DMRs may be distorted by pre-natal confounders. For the remaining six DMRs, there was no evidence for directional concordance between any of the two comparisons, suggesting that the association between breastfeeding and methylation at age 7 in the CpGs in those DMRs may be transient or false-positives. A sensitivity analysis considering only those CpGs that achieved P<0.05 in at least one time point corroborated the strongest directional consistency between 7 and 15-17 years observed for the four aforementioned DMRs, except the 8:409905-410098; importantly, this analysis involved only 3 CpGs for this DMR (Supplementary Table 4). Moreover, a fifth DMR – 19:365914-366989 – was identified in this analysis, suggesting that CpGs with weak associations could have diluted the association in the analysis considering all CpGs in the DMR.

**Table 3.** Association between peripheral blood DNA methylation at different ages at each DMR and ever breastfeeding.

| DMR<br>(Chr:Start-End<sup>a</sup>) | At birth |  |  | 7 years |  |  | 15-17 years |  |  |
| --- | --- | --- | --- | --- | --- | --- | --- | --- | --- |
| | $\beta$ | SE | P-value | $\beta$ | SE | P-value | $\beta$ | SE | P-value |
| 5:97,867-98,797 | 0.30 | 0.21 | 0.146 | 0.43 | 0.21 | 0.043 | 0.30 | 0.21 | 0.158 |
| 19:365,914-366,989 | -0.01 | 0.34 | 0.975 | 0.05 | 0.34 | 0.881 | -0.04 | 0.35 | 0.897 |
| 18:106,178-106,850 | -0.08 | 0.77 | 0.913 | 0.14 | 0.75 | 0.855 | 0.23 | 0.77 | 0.767 |
| 1:425,524-426,297 | 0.26 | 0.62 | 0.673 | 0.33 | 0.61 | 0.590 | 0.16 | 0.62 | 0.800 |
| 9:91,296-92,146 | -0.10 | 0.33 | 0.759 | -0.18 | 0.33 | 0.578 | -0.10 | 0.34 | 0.755 |
| 17:222,498-222,991 | -0.01 | 0.37 | 0.983 | 0.00 | 0.36 | 0.994 | -0.04 | 0.36 | 0.913 |
| 4:136,643-137,027 | -0.03 | 0.41 | 0.951 | -0.37 | 0.38 | 0.324 | -0.31 | 0.41 | 0.448 |
| 22:255,590-256,045 | 0.40 | 0.71 | 0.577 | 1.18 | 0.70 | 0.095 | 1.06 | 0.71 | 0.136 |
| 4:33,482-33,808 | 0.13 | 2.05 | 0.950 | 0.06 | 2.00 | 0.978 | 0.08 | 2.04 | 0.967 |
| 8:409,905-410,098 | 0.82 | 1.31 | 0.530 | 1.05 | 1.32 | 0.425 | 1.04 | 1.32 | 0.433 |
| 1:224,191-225,190 | 0.03 | 0.45 | 0.940 | -0.03 | 0.44 | 0.951 | -0.03 | 0.45 | 0.948 |
| 9:61,093-61,964 | -0.39 | 0.50 | 0.432 | -0.44 | 0.49 | 0.369 | -0.39 | 0.50 | 0.435 |

Regression coefficients ( $\beta$ ) are average percent point differences in DNA methylation averaged across CpGs that belong to the DMR. This analysis was performed using linear mixed models to account for the correlation between CpGs in the same DMR.
<sup>a</sup>Human Genome Assembly GRCh37.
Chr: Chromosome. DMR: differentially methylated region. SE: standard error.

**Table 4.** Directional concordance between time points for each individual CpG belonging to the same DMR.

| DMR<br>(Chr:Start-End<sup>a</sup>) | Number<br>of CpGs | At birth and 7 years |  | 7 years and 15-17 years |  |
| --- | --- | --- | --- | --- | --- |
|  |  | Concordance | P-value | Concordance | P-value |
| 5:97,867-98,797 | 275 | 66.2 | $8.7 \times 10^{-8}$ | 69.1 | $2.2 \times 10^{-10}$ |
| 19:365,914-366,989 | 205 | 47.8 | 0.576 | 54.1 | 0.264 |
| 18:106,178-106,850 | 18 | 72.2 | 0.096 | 83.3 | 0.008 |
| 1:425,524-426,297 | 64 | 68.8 | 0.004 | 56.3 | 0.382 |
| 9:91,296-92,146 | 185 | 54.1 | 0.303 | 58.4 | 0.027 |
| 17:222,498-222,991 | 140 | 55.7 | 0.205 | 49.3 | 0.933 |
| 4:136,643-137,027 | 13 | 69.2 | 0.267 | 61.5 | 0.581 |
| 22:255,590-256,045 | 30 | 63.3 | 0.200 | 83.3 | $3.3 \times 10^{-4}$ |
| 4:33,482-33,808 | 5 | 60.0 | 0.999 | 60.0 | 0.999 |
| 8:409,905-410,098 | 7 | 85.7 | 0.125 | 100.0 | 0.016 |
| 1:224,191-225,190 | 129 | 57.4 | 0.113 | 47.3 | 0.597 |
| 9:61,093-61,964 | 91 | 57.1 | 0.208 | 56.0 | 0.294 |

Concordance is shown in %. The analyses were performed using a sign test.
<sup>a</sup>Human Genome Assembly GRCh37.
Chr: Chromosome. DMR: differentially methylated region.

## Discussion

In this epigenome-wide association study, having ever been breastfed was associated with peripheral blood methylation in the cg11414913 CpG at ages 7 and 15-17 years, but not at birth. There was suggestive evidence of association between ever been breastfed and age 7 methylation in six additional CpGs, with one – the cg05800082 CpG – also presenting a directionally consistent (although attenuated) point estimate at age 15, but not at birth. Moreover, 12 DMRs were identified at age 7, and three of them presented evidence of directional concordance between ages 7 and 15-17, but not between birth and age 7, in all sensitivity analyses. Our quantile-quantile plots indicated that the associational effect estimates between ever breastfeeding and peripheral blood DNA methylation are generally small. None of our analyses supported a dose-response relationship between breastfeeding and peripheral blood DNA methylation, but were consistent with an effect that depends on whether or not the child was ever breastfed.

The epidemiological literature on breastfeeding and health focuses on well-established effects against infectious diseases, as well as on potential impact on intelligence, obesity and diabetes, among other outcomes [1]. In the present analyses, only one site where methylation differences were detected could be involved in the above traits. Specifically, an online search into the biological role of the genes whose methylation was associated with breastfeeding (see Supplementary info) showed an effect on the *DST* gene, which is expressed in many tissues, including the brain, where it encodes isoforms of cytoskeletal linker proteins anchor neural intermediate filaments to the actin cytoskeleton. Other sites affected were not located on genes with known function or were in genes expressed in other tissues such as testis. This may be due to analysing a surrogate tissue, limited statistical power to detect more CpGs, and limited knowledge about the health effects of the methylation sites that were detected. Moreover, the effects of breastfeeding on health and development may be mediated through other epigenetic processes, such as non-coding RNAs [35, 36], as well as a host of mechanisms other than epigenetics, including provision of nutrients (e.g., pre-formed long-chain polyunsaturated fatty acids, which is a plausible mediator of the benefits on IQ [37]), antibodies and other immunoactive compounds, antimicrobials, and important effects on the gut microbiome [1].

One of the strengths of this study is that longitudinal measures of DNA methylation allowed not only identifying regions of the methylome associated with breastfeeding, but also assessing if those associations persist until adolescence. Dense phenotyping and genotyping of study participants allowed controlling for several covariates, which were selected using a conceptual model defined *a priori*. Moreover, DNA methylation data at birth was used to rule out associations likely driven by residual confounding due to pre-natal factors. To unravel the possibility of residual confounding by maternal smoking, we checked the overlap between the suggestive CpG sites from our breastfeeding EWAS and the largest maternal smoking EWAS [38], and found that none of the sites were amongst the 6073 sites that were associated with maternal smoking during pregnancy, suggesting that it is unlikely that the associations were driven by residual confounding by maternal smoking. However, residual confounding cannot be fully discarded due to missing confounders (including post-birth factors that may affect breastfeeding quality and duration) measurement error and model misspecification. Therefore, triangulating our findings with those from future studies using designs prone to different potential sources of bias will be important to disentangle causality [39].

In addition to the possibility of residual confounding, another weakness of our study is that our exposure variable was ill-defined. Due to sample size constraints and limitations of self-reported data, it was not possible to use more refined definitions of breastfeeding. Indeed, our main results were related to the binary categorisation, which includes, in the “breastfed” group, highly heterogeneous individuals regarding breastfeeding quality, duration, type of foods given concurrently with breastmilk (for individuals that were non-exclusively breastfeed) and after weaning, among other factors. Similarly, the “non-breastfed” group potentially includes individuals that received many different types of foods. This heterogeneity is likely to influence the results in ways that are rather difficult to predict, and limits the external validity of our findings.

It should also be noted that our study was restricted to peripheral blood. As we discussed elsewhere [18], DNA methylation in blood is unlikely to be a good proxy of DNA methylation in other tissues, such as the brain [40, 41, 42], thus limiting the capacity of any breastfeeding EWAS using peripheral blood to inform DNA methylation patterns in the target tissue [11, 43] – in this example, when assessing if the association between breastfeeding and IQ has a component related to methylation. This may also limit the capacity to identify true signals. However, DNA methylation studies in surrogate tissues are important. These are frequently the only viable alternative in large epidemiological studies, also being able to provide useful information on the range of potential effects of the exposure of interest on DNA methylation, which may then guide future, specific studies such as *in vitro* studies in cells and *in vivo* studies in animal models [18].

Another important limitation is that we did not perform a formal replication of our results. However, the fact that some hits (both in the CpG and DMR analysis) at age 7 years did not present evidence of association at age 15-17 years indicates that inflation of type-I error due to multiple-testing alone was not sufficient for a hit in one age to also present evidence of association in other ages. Therefore, CpGs and DMRs that presented evidence of persistent associations are less likely to be a sole product of multiple testing. However, this reasoning is less clear for transient associations, which could be truly transient effects or merely false-positives that do not carry over to adolescence. Although persistent associations are likely to be more robust from a methodological perspective in our study, this does not mean that transient effects are irrelevant. For example, they could trigger the actual processes related to long-term effects (e.g., influences on brain development and IQ in adulthood). Moreover, in our context transient effects mean that associations observed at the age of 7 years did not persist until adolescence, but associations at age 7 would already be persistent effects of breastfeeding. Finally, it is important to consider the loss of individuals to missing data. About 30% of ARIES data were removed due to missing exposure, covariate or outcome data, which reduces the power to find CpG sites related to breastfeeding. Methods for multiple imputation in methylation data [44] are at an early stage and therefore were not used here, but in future these methods will be crucial to maximise the power of an EWAS.

## Conclusions

This study provides important insights into the magnitude and persistence of the association between breastfeeding and peripheral blood DNA methylation. Rather than providing definitive answers on their own, our results will serve to motivate future studies using different designs to improve causal inference, as well as consortium-based efforts – examples of which are already available in the epigenetic epidemiology literature [38, 45] – to achieve sample sizes large enough to both improve power and allow replication. Such future efforts will complement and expand our findings by providing robust evidence on the potential effects of breastfeeding on DNA methylation, which may contribute to understand the biological basis of long-term associations between breastfeeding and health and human capital outcomes, and potentially also reveal new biological aspects of breastfeeding.

## List of abbreviations

ALSPAC: Avon Longitudinal Study of Parents and Children.
ARIES: Accessible Resource for Integrated Epigenomic Studies.
DAG: directed acyclic graoph.
DMR: differentially methylated region.
DNA: deoxyribonucleic acid.
EWAS: epigenome-wide association study.
IQ: intelligence quotient.

## Declarations

### Ethics approval and consent to participate

Only individuals who gave informed and written consent were enrolled in ALSPAC. Ethical approval for the study was obtained from the ALSPAC Ethics and Law Committee and the Local Research Ethics Committees.

### Consent for publication

Not applicable.

### Availability of data and material

The datasets analysed during the current study are not publicly available due to them containing information that could compromise research participant privacy/consent, but they are available on request to the Executive.

### Authors’ contributions

Study conception and design: CGV, CR, DC, FPH and GDS.

Data acquisition: CR and GDS.

Data analysis and interpretation: AJS, DC and FPH.

Manuscript writing: DC and FPH.

Critical revision of the manuscript: AJS, CGV, CR, GDS.

All authors read and approved the final manuscript.

## Acknowledgements

We are extremely grateful to all the families who took part in this study, the midwives for their help in recruiting them, and the whole ALSPAC team, which includes interviewers, computer and laboratory technicians, clerical workers, research scientists, volunteers, managers, receptionists, and nurses.

