## Supplementary text and tables for "Association between breastfeeding and DNA methylation over the life course: findings from the Avon Longitudinal Study of Parents and Children (ALSPAC)"

**Biological characterisation of the CpGs and DMRs**

**Methods**

The nearest gene to each CpG site was extracted from the annotation file provided by Illumina. We used the UCSC Genome Browser (<https://genome.ucsc.edu/cgi-bin/hgGateway>; GRCh37/hg19 Assembly) where no genes were available, and to identify other biological features – focusing on DNase I hypersensitivity, presence of binding sites of transcription factors, and conservation among vertebrates – of the regions containing the identified CpGs and DMRs. Features of identified genes (and encoded proteins) were extracted from GeneCards®: The Human Gene Database (<http://www.genecards.org/>) and from GeneEntrez (<https://www.ncbi.nlm.nih.gov/gene>). Linked diseases were identified using the Online Mendelian Inheritance in Man (OMIM) database (<https://www.ncbi.nlm.nih.gov/omim/>).

**Results**

cg11414913, which presented the most robust statistical evidence of association with breastfeeding, is located in an intergenic region, with the nearest gene being the *TTC34* gene. This gene is overexpressed in the testis, but largely unknown regarding its biological roles, although there is some indication of a relation with multiple sclerosis and lung cancer. The region around this CpG is highly conserved among vertebrates and contains a 249 bp region (which includes the CpG) that presents DNase I hypersensitivity (which is related to more transcriptional activity) in six cell/tissue types, including lung carcinoma, prostate adenocarcinoma and pancreatic islets. cg05800082, which presented some evidence of persistent association with breastfeeding, is located within the *DST* gene, which is expressed in many tissues, including skin and brain. This gene encodes isoforms of cytoskeletal linker proteins that present tissue-specificity regarding expression and function: while some isoforms expressed in epithelial tissues anchor keratin-containing intermediate filaments to hemidesmosomes, other isoforms – mainly expressed in neural and muscle tissue – anchor neural intermediate filaments to the actin cytoskeleton. Mutations in the *DST* gene have also been implicated in neuronal and skin disorders. Moreover, the region spanning this CpG presents DNase I hypersensitivity in 5 cell/tissue types and enrichment of the H3K27Ac histone mark, which is also related to enhanced transcription.

Regarding DMRs, the 18:106,178-106,850 region is located within the *DUX4* gene, which encodes a transcriptional activator of *PITX1*, and is linked to autosomal dominant facioscapulohumeral muscular dystrophy (FSHD). It is expressed in the testis, and in muscle tissues of FSHD patients. The 9:91,296-92,146 region is located 1,719 bp away from the *PGM5P3-AS1* gene, which encodes a non-coding RNA of unknown function. The 22:255,590-25,6045 region did not present any obvious important biological feature in a 100,000 bp window centred at the DMR. Two additional DMRs presented weaker evidence of a persistent association with breastfeeding. One was the 8:409,905-410,098 region located in the *FBXO25* gene, which encodes a protein that is overexpressed in the testis and belong to the family of F-box proteins, which are components of a ubiquitin protein ligase complex. The second was the 19:365,914-366,989 region located in the *THEG* gene, which encodes a nuclear protein specifically in the nucleus of haploid male germ cells, with a possible role in spermatogenesis.

**Supplementary Tables**

**Supplementary Table 1.** Description of the individuals included in the main analysis, compared to all ARIES participants, restricting to those with age 7 methylation data available.

| **Variable** | **Statistic/category^a^** | **All ARIES participants (n=995)** | **Participants included in this study (n=702)** |
| --- | --- | --- | --- |
| Maternal education | CSE | 8.9% | 7.2% |
| at birth | Vocational education | 7.4% | 6.0% |
|  | GCE Ordinary level | 34.3% | 33.8% |
|  | GCE Advanced level | 29.1% | 29.9% |
|  | Degree | 20.3% | 23.1% |
| Maternal age at birth (years) | Mean (SD) | 29.5 (4.4) | 30.0 (4.4) |
| Parity | 0 | 46.5% | 45.7% |
|  | 1 | 36.9% | 37.5% |
|  | 2 | 12.7% | 13.4% |
|  | ≥3 | 3.9% | 3.4% |
| Maternal smoking | Never | 86.3% | 87.7% |
| in relation to | Before | 3.7% | 4.0% |
| pregnancy | During | 10.0% | 8.3% |
| Folic acid | No | 75.9% | 75.9% |
| supplementation | Yes | 24.1% | 24.1% |
| Caesarean section | No | 90.4% | 90.2% |
|  | Yes | 9.6% | 9.8% |
| Birthweight (g) | Mean (SD) | 3487 (486) | 3490 (476) |
| Sex | Male | 48.9% | 49.1% |
|  | Female | 51.1% | 50.9% |
| Ethnicity | European | 97.0% | >99.0% |
|  | Other | 3.0% | <1.0%^b^ |
| Breastfeeding duration | 0 | 11.1% | 10.4% |
| (months) | 0.1-3 | 32.0% | 31.0% |
|  | 3.1-6 | 16.2% | 16.2% |
|  | 6.1-12 | 27.6% | 28.2% |
|  | >12 | 13.1% | 14.2% |

^a^Mean and SD for continuous variables, and each category (for which proportions are shown) for categorical variables.

^b^Masked for disclosure purposes.

CSE: Certificate of Secondary Education. GCE: General Certificate of Education. SD: standard deviation.

**Supplementary Table 2**. Description of the CpGs that presented at least suggestive evidence of association with ever breastfeeding in the fully-adjusted analysis at age 7.

| **CpG** | **Chromosome: position (bp)^a^** | **Nearest gene** | **Distance (bp) to nearest gene** |
| --- | --- | --- | --- |
| cg11414913 | 1:2,799,662 | *TTC34* | 93,432 |
| cg00234095 | 17:39,440,474 | *KRTAP9-7* | 8,015 |
| cg04722177 | 19:39,737,768 | *IFNL4* | Intragenic |
| cg03945777 | 7:157,514,049 | *PTPRN2* | Intragenic |
| cg17052885 | 17:78,896,012 | *RPTOR* | Intragenic |
| cg05800082 | 6:56,508,429 | *DST* | Intragenic |
| cg24134845 | 10:100,992,149 | *HPSE2* | Intragenic |

^a^Human Genome Assembly GRCh37.

bp: base pairs.

**Supplementary Table 3.** Differentially methylated regions (DMR) in peripheral blood at age 7 according to ever breastfeeding.

| **DMR**  **(Chr:Start-End^a^)** | **Number**  **of CpGs** | **P-value** | **Nearest gene** | **Distance (bp) to**  **nearest gene** |
| --- | --- | --- | --- | --- |
| 5:97,867-98,797 | 275 | 3.2×10^-6^ | *PLEKHG4B* | Intragenic |
| 19:365,914-366,989 | 205 | 9.7×10^-5^ | *THEG* | Intragenic |
| 18:106,178-106,850 | 18 | 0.001 | *DUX4* | Intragenic |
| 1:425,524-426,297 | 64 | 0.002 | *BC036251* | 4,458 |
| 9:91,296-92,146 | 185 | 0.003 | *PGM5P3-AS1* | 1,719 |
| 17:222,498-222,991 | 140 | 0.003 | *RPH3AL* | 19,865 |
| 4:136,643-137,027 | 13 | 0.007 | *ZNF595*/*ZNF718* | Intragenic |
| 22:255,590-256,045 | 30 | 0.012 | *AK022914* | 15,894,215 |
| 4:33,482-33,808 | 5 | 0.019 | *ZNF595*/*ZNF718* | 19,419 |
| 8:409,905-410,098 | 7 | 0.025 | *FBXO25* | Intragenic |
| 1:224,191-225,190 | 129 | 0.045 | *LOC729737* | 83,625 |
| 9:61,093-61,964 | 91 | 0.046 | *AY343892* | 10,734 |

^a^Human Genome Assembly GRCh37.

^b^No gene within a 100,000 bp window centred at this region.

Chr: chromosome. bp: base pairs.

**Supplementary Table 4.** Directional concordance (in %) between time points for each individual CpG belonging to the same differentially methylated region (DMR). Only CpGs that achieved P<0.05 in at least one time point were considered.

| **DMR** | **Number** | **At birth and 7 years** | | **7 years and 15-17 years** | |
| --- | --- | --- | --- | --- | --- |
| **(Chr:Start-End^a^)** | **of CpGs** | **Concordance** | **P-value** | **Concordance** | **P-value** |
| 5:97,867-98,797 | 69 | 72.5 | 2.4×10^-4^ | 85.5 | 1.4×10^-9^ |
| 19:365,914-366,989 | 38 | 52.6 | 0.871 | 68.4 | 0.034 |
| 18:106,178-106,850 | 8 | 75.0 | 0.289 | 100.0 | 0.008 |
| 1:425,524-426,297 | 15 | 80.0 | 0.035 | 73.3 | 0.118 |
| 9:91,296-92,146 | 38 | 63.2 | 0.143 | 71.1 | 0.014 |
| 17:222,498-222,991 | 22 | 45.5 | 0.832 | 50.0 | 0.999 |
| 4:136,643-137,027 | 3 | 100.0 | 0.250 | 66.7 | 0.999 |
| 22:255,590-256,045 | 16 | 68.8 | 0.210 | 93.8 | 0.001 |
| 4:33,482-33,808 | 2 | 50.0 | 0.999 | 0.0 | 0.500 |
| 8:409,905-410,098 | 3 | 100.0 | 0.250 | 100.0 | 0.250 |
| 1:224,191-225,190 | 23 | 39.1 | 0.405 | 56.5 | 0.678 |
| 9:61,093-61,964 | 24 | 75.0 | 0.023 | 66.7 | 0.152 |

^a^Human Genome Assembly GRCh37.

Chr: Chromosome.
